# Predicting drug concentrations in PDMS microfluidic organ chips

**DOI:** 10.1101/2021.05.01.442248

**Authors:** Jennifer Grant, Alican Özkan, Crystal Oh, Gautam Mahajan, Rachelle Prantil-Baun, Donald E. Ingber

## Abstract

Microfluidic organ-on-a-chip (Organ Chip) cell culture devices are often fabricated using polydimethylsiloxane (PDMS) because it is biocompatible, transparent, elastomeric, and oxygen permeable; however, hydrophobic small molecules can absorb to PDMS, which makes it challenging to predict drug responses. Here, we describe a combined simulation and experimental approach to predict the spatial and temporal concentration profile of a drug under continuous dosing in a PDMS Organ Chip containing two parallel channels separated by a porous membrane that is lined with cultured cells, without prior knowledge of its log P value. First, a three-dimensional finite element simulation of drug loss into the chip was developed that incorporates absorption, adsorption, convection, and diffusion, which predicts changes in drug levels over time and space as a function of potential PDMS diffusion coefficients and log P values. By then experimentally measuring the diffusivity of the compound in PDMS and determining its partition coefficient through mass spectrometric analysis of the drug concentration in the channel outflow, it is possible to estimate the effective log P range of the compound. The diffusion and partition coefficients were experimentally derived for the antimalarial drug and potential SARS-CoV-2 therapeutic, amodiaquine, and incorporated into the model to quantitatively predict the drug-specific concentration profile over time measured in human Lung Airway Chips lined with bronchial epithelium interfaced with pulmonary microvascular endothelium. The same strategy can be applied to any device geometry, surface treatment, or *in vitro* microfluidic model to predict the spatial and temporal gradient of a drug in 3D without prior knowledge of the partition coefficient or the rate of diffusion in PDMS. Thus, this approach may expand the use of PDMS Organ Chip devices for various forms of drug testing.

## INTRODUCTION

Organs-on-chips (Organ Chips) are microfluidic culture devices lined by living human cells that reconstitute human organ-level structures as well as physiology and pathophysiology. Their ability to model many features of a functional human organ, such as the response to viral infection,^1,2^ bacterial infection,^3,4^ disease states,^5–9^ and clinically relevant drug exposure pharmacokinetic (PK) profiles,^10^ make Organ Chips particularly useful for drug development and toxicity studies. Organ Chips are often fabricated out of poly-dimethylsiloxane (PDMS) because it is an oxygen permeable, transparent, biocompatible, and elastomeric polymer.^3,11^ However, PDMS can strongly adsorb some hydrophobic small molecules from solution as well as accumulate these molecules into its bulk volume. Together, adsorption and absorption can decrease the effective dosing concentration, making it difficult to predict actual drug concentrations experienced by the cells. Thus, development of a method to overcome this limitation could benefit the entire Organ Chip field.

The logarithm of the octanol-water partition coefficient (log P) is frequently used to approximate the extent of PDMS absorption, where compounds with high log P have a greater tendency to bind to or be absorbed by PDMS. Partition coefficients have been experimentally determined using analytical techniques such as fluorescence,^12^ UV-vis,^13^ or IR spectroscopy,^13^ and computationally predicted with quantitative structure-activity relationship (QSAR)^14,15^ or molecular dynamics (MD)^16^ models. However, the partition coefficient alone cannot predict the extent of binding, and past work suggests that it should be combined with the topological polar surface area (TPSA)^12,17^ or the number of H-bond donors^13^ to improve the accuracy of bioavailability predictions in PDMS-based microfluidic devices. Although physiochemical descriptors are useful for describing the extent of compound loss into PDMS devices, they cannot be used to quantitatively predict drug loss because it is a dynamic process that depends strongly on a variety of factors including absorption, convection, diffusion, medium composition, dosing time, cell type, and surface treatment, in addition to adsorption.

A more comprehensive understanding of drug loss can be obtained by combining computational models and experimental data acquired directly from the PDMS Organ Chip devices used for *in vitro* culture. A one-dimensional (1D) simulation model has been developed that predicts drug loss in a PDMS device by combining computational fluid dynamics (CFD) with drug-PDMS binding kinetics, which suggested that drug loss can be minimized by increasing the flow rate, decreasing channel height, or decreasing channel width.^13^ This approach demonstrated the potential to estimate the concentration of the drug in a simple single-channel microfluidic culture device by combining experimentally derived constants with computational simulations of physical processes. However, it does not provide a way to predict how drug concentrations vary over time and space within microfluidic Organ Chip devices that contain two parallel channels separated by a membrane lined by living cells, which have been shown to faithfully recapitulate the physiology and pathophysiology of many different human organs.^10,18^ Two-channel microfluidic Organ Chips can faithfully recapitulate dynamic pharmacokinetic (PK) profiles that are observed *in vivo* because the drug is perfused through endothelium-lined vascular channels and physiologically relevant tissue-tissue interactions permit analysis of clinically relevant drug fluxes between compartments. We have previously shown that PK parameters can be estimated in a vascularized human bone marrow-on-a-chip (BM chip) by fitting a three-compartment PK model to LC-MS/MS-quantified drug concentrations from the chip outflows.^19^ It is possible to increase the accuracy of the PK model and *in vivo* extrapolations by accounting for drug loss due to PDMS absorption. A PK model of a Human-Body-on-Chips system incorporated drug loss due to PDMS absorption by performing mass spectrometric analysis of the drug concentration in the outflows of blank chips and incorporating the fractional loss into the model.^10^ Although this model quantitatively predicted PK parameters for orally and intravenously administered drugs, it cannot be used to accurately predict the spatial and temporal concentration of the drug inside the chip throughout the dosing period because the model did not consider diffusion of the drug into bulk PDMS and it simplified the geometry of the Organ Chip into two-dimensional (2D) domains. Additionally, the loss due to PDMS absorption was performed by mass spectrometric analysis of blank chips without cultured cells, which can bias the extent of absorption. Thus, there is a clear need for a more comprehensive model that simulates three-dimensional (3D) drug concentrations in chips lined with cultured cells and incorporates all physical processes that contribute to drug loss. This is necessary to quantitatively predict drug concentrations inside the chip throughout an exposure period and thereby guide dosing strategies, evaluate toxicities, and analyze pharmacokinetic properties.

Here, we describe a combined experimental and computational approach that predicts spatial and temporal drug concentration profiles in 3D under continuous dosing in two-channel, microfluidic, Organ Chips lined with cultured living cells. This strategy involves the development of a finite element simulation of drug absorption in a PDMS Organ Chip as well as experimental quantification of the diffusion and partition coefficients of the drug. The computational model is then used to predict drug concentrations that the cells experience at any time within the microfluidic channels of the chip. We applied this method towards predicting levels of the antimalarial drug amodiaquine when administered continuously under flow in human Lung Airway Chips, which recently resulted in identification of this drug as a potential SARS-CoV-2 therapeutic.^1^ This strategy can predict compound loss due to PDMS absorption in any device composition, and thus, it should help to improve experimental design and analysis of dose-response efficacy and toxicity studies in PDMS Organ Chips.

## RESULTS

### Simulating Drug Concentration in Two-Channel PDMS Organ Chips

The mass transport of a drug in a microfluidic Organ Chip is controlled by diffusion, convection, dissolution, dose, device geometry, temperature, and pressure.^20^ We carried out 3D simulations that incorporate each of these factors to understand the drug distribution in the microfluidic channels. The Organ Chip we model is a commercially available PDMS microfluidic culture device that contains two parallel fluidic channels separated by a porous PDMS membrane, which is lined by living epithelium on one side and endothelium on the other (**Fig. 1a**). A SOLIDWORKS rendering of the PDMS Organ Chip was imported into COMSOL Multiphysics software and fluidic domains were assigned to the apical and basal channels (**Fig 1b**). The height of the epithelium and endothelium (3.9 μm and 1 μm, respectively) was experimentally determined from microscopic cross-sections of human Lung Large Airway Chips lined by primary human bronchial epithelium interfaced with primary human pulmonary microvascular endothelium and incorporated into the total height of the central membrane; the membrane pores that represent 3% of the total membrane area were removed to increase computational efficiency.

**Figure 1.**
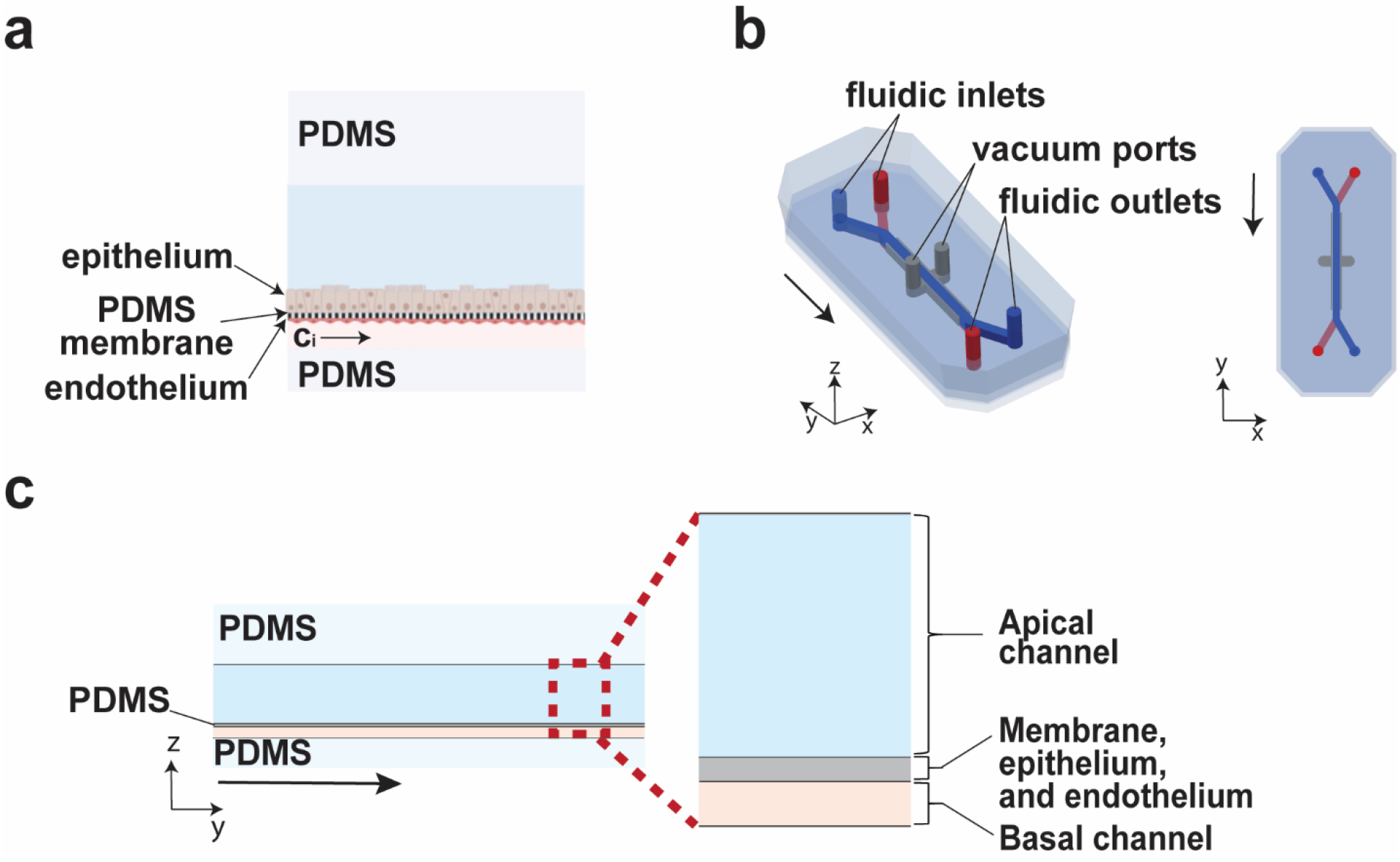
Design of a two-channel human Organ Chip. **a**) Side-view schematic of the microfluidic chip with epithelium lining the top of the horizontal membrane that separates the upper and lower channels, interfaced with endothelium that is cultured on its basal surface. Drug is dosed at a concentration (c_i_) in the endothelium-lined channel under continuous perfusion to mimic systemic distribution after oral administration (image generated with BioRender). **b**) The Organ Chip geometry simulated in COMSOL. The Organ Chip has two central fluidic channels (blue apical channel and red basal channel) that are separated by a porous membrane. The fluidic channels are positioned between two full height vacuum chambers (grey), although they were not used to apply cyclic strain during the culture of these large airway cells. The two channels and vacuum chambers all have inlets and outlets that vertically rise to the upper surface of the chip. **c**) Side-view image of the simulated Organ Chip with the inset showing the apical channel, the central membrane lined with epithelium on its upper surface and endothelium on its underside, and the basal fluidic channel. Arrows indicate the flow direction.

We used the human Lung Airway Chip as a starting point to develop this modeling approach because we had found that the antimalarial drug, amodiaquine, significantly inhibits infection of the epithelium by a pseudotyped SARS-CoV-2 virus when flowed through the vascular channel of the device at a clinically relevant dose of 1.24 μM (the maximum concentration or C_max_ of amodiaquine in human blood), and its ability to inhibit infections by native SARS-CoV-2 was confirmed *in vitro* and in animal studies.^1^ Amodiaquine is known to participate in hydrophobic interactions with lipids and proteins,^21,22^ and thus, there was a good chance that it might adsorb to the walls of PDMS channels. So, we wanted to determine the effective drug concentration that the cells were exposed to in this microfluidic chip under the experimental conditions we utilized; however, we did not know the log P for this drug in the Organ Chip environment with cultured cells and extracellular matrix. Thus, to fully characterize the concentration of amodiaquine in the Lung Airway Chip, we set out to develop a combined experimental and computational approach that predicts the concentration of the drug in the chip over time.

A drug must first adsorb to the channel wall before it enters into the bulk PDMS. Dissolution of a drug from the cell culture medium onto the PDMS wall is described using the partition coefficient (P):

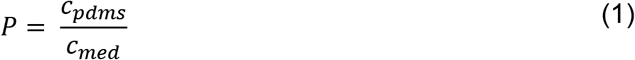

where *c*_*pdms*_ is the concentration of drug in PDMS and *c*_*med*_ is the concentration of drug in cell culture medium. The partition coefficient was implemented for all medium-PDMS interfaces in the Organ Chip geometry. After adsorption onto the wall surface, the drug diffuses into the bulk polymer down a concentration gradient at a rate described by Fick’s second law of diffusion. In our model, transport of this small hydrophobic molecule through the porous membrane and living tissue-tissue interface was assumed to be passive and simplified to follow Fick’s second law. We simulated continuous perfusion of a small molecule with a size similar to that of amodiaquine (~400 Da) at 1.24 μM, under continuous flow (60 μL/h) through the basal vascular channel to mimic systemic dosing after oral administration; the drug was also maintained under static conditions in the apical channel, as carried out in the published study.^1^ A time-dependent simulation was performed for 12 h with the temperature, flow rate, atmospheric pressure, diffusion coefficient of drug in medium, diffusion coefficient of drug in PDMS, and partition coefficient being held constant.

First, we explored the effect of varying the partition coefficient on the distribution of the small molecule when flowed through the basal channel of the Organ Chip for 12 h. The simulation was performed with P= 100 and 400 (log P= 2 and 2.6, respectively) and D_pdms_ = 1 × 10^−13^ m^2^/s, which is the approximated diffusion coefficient for a compound with molecular weight ~400 Da.^23–25^ 3D surface heat maps of the drug concentration in the apical and basal channel show how the drug accumulates in both fluidic channels throughout the simulated dosing period (**Fig. 2a**). 2D heat maps of a vertical cross section through the center of the chip in the xz axis show that absorption to the channel wall generates a concentration gradient that radially extends outwards from the center of the basal channel with lower concentrations along its lateral sidewalls (**Fig. 2b**). The concentration gradient in the basal channel is anisotropic because the 90° corners generate a non-uniform velocity distribution (**Fig. S1**). These models reveal that the drug with P= 400 takes more time to saturate the basal channel because it rapidly absorbs into the channel walls. At both simulated P values, drug enters the membrane from the basal channel and begins to diffuse into the apical channel medium (**Fig. 2b** and **Fig. S2**). 2D heat maps through the center of the basal channel along the xy plane show that a concentration gradient also exists along the fluidic stream because adsorption depletes the drug from the medium and lowers the downstream concentration along the length of the channel (**Fig. 2c**). Together, these data emphasize that drug absorption and adsorption are spatially and temporally dependent processes and both influence the concentration of drug dose exposure experienced in this *in vitro* chip model. Thus, applying a single constant to describe drug loss in an Organ Chip will not capture these dynamic concentration distributions and may inaccurately describe the dosing window.

**Figure 2.**
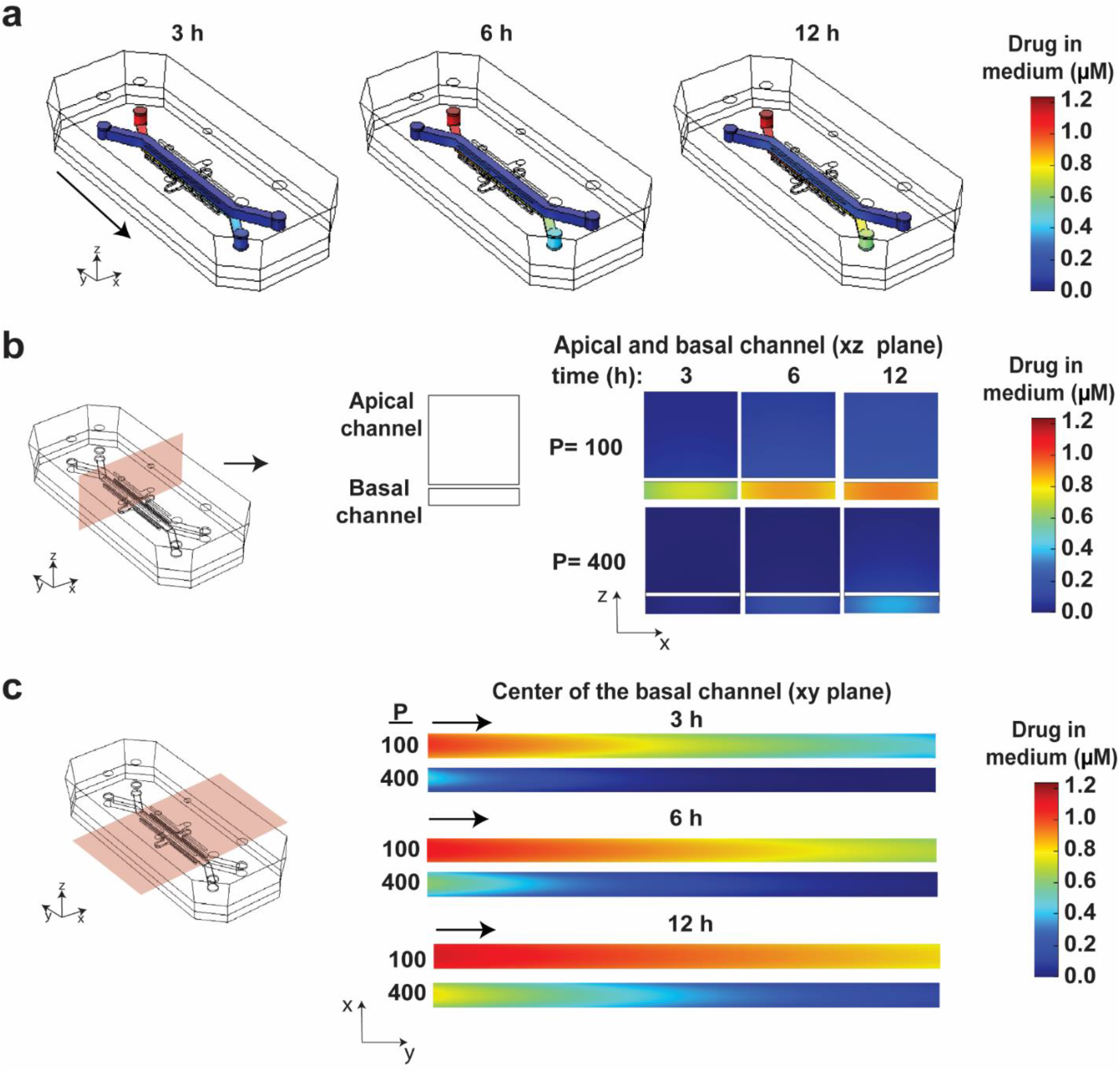
Heat maps of the drug concentration in the chip over time show that loss due to adsorption and absorption is a spatially and temporally dynamic process. **a**) 3D surface heat maps of the concentration of drug in the fluidic channels of the human Lung Airway Chip viewed from above after 3, 6, and 12 h with P=100 and D_pdms_ = 1 × 10^−13^ m^2^/s. Drug is continuously flowed into the basal channel and the apical channel is held static. Arrow indicates the flow direction. **b**) 2D heat maps of drug concentration through a vertical cross section in the center of the chip along the xz plane at 3, 6, and 12 h for P=100 and 400, and D_pdms_ = 1 × 10^−13^ m^2^/s. The heat maps show the concentration of drug in the apical and basal channel medium in the vertical chip cross section. The drug concentration radially extends outwards from the center of the channel with the lowest concentrations near the lateral side walls and begins to enter the apical channel fluid over time. Drugs with lower P values require less time to reach the center of the channel. **c**) 2D heat maps of drug concentration through a horizontal cross section in the center of the basal channel along the xy plane at 3, 6, and 12 h for P=100 and 400 and D_pdms_ = 1 × 10^−13^ m^2^/s. Arrows indicate the flow direction.

Next, we explored the effect of D_pdms_ and P on the concentration of the drug in the basal channel when continuously dosed at 1.24 μM under 60 μL/h flow for 12 h. We varied D_pdms_ from 1 × 10^−12^ m^2^/s to 1 × 10^−15^ m^2^/s and observed that compounds with low PDMS diffusivity (D_pdms_ ≤ 1 × 10^−13^ m^2^/s) require less time to saturate in the fluidic channel because the compound accumulates in bulk PDMS near the PDMS-medium interface (**Fig. 3a**). Conversely, more drug loss is observed for compounds with high PDMS diffusivity (D_pdms_ > 1 × 10^−13^ m^2^/s) because the compound travels down its concentration gradient faster and does not accumulate near the wall. We also varied P from 25-600 while keeping D_pdms_ constant at 1 × 10^−13^ m^2^/s and observed that drugs with higher P values (P>200 = log P>2.3) show significant depletion in the basal channel (**Fig. 3b**). As expected, drugs with high partition coefficients adsorb more rapidly to the channel walls and deplete from the medium at a higher rate. The simulations emphasize that *in vitro* models with the same flow rate and dosing concentration can experience dramatically different compound loss depending on the value of P and D_pdms_. Our work complements past studies which concluded that drug loss in PDMS can be significant for log P > 2.7, but total loss also depends on diffusivity, convection, and dosing time.^26^ Therefore, drug loss should be simulated and evaluated under the same conditions as used in the *in vitro* model of interest.

**Figure 3.**
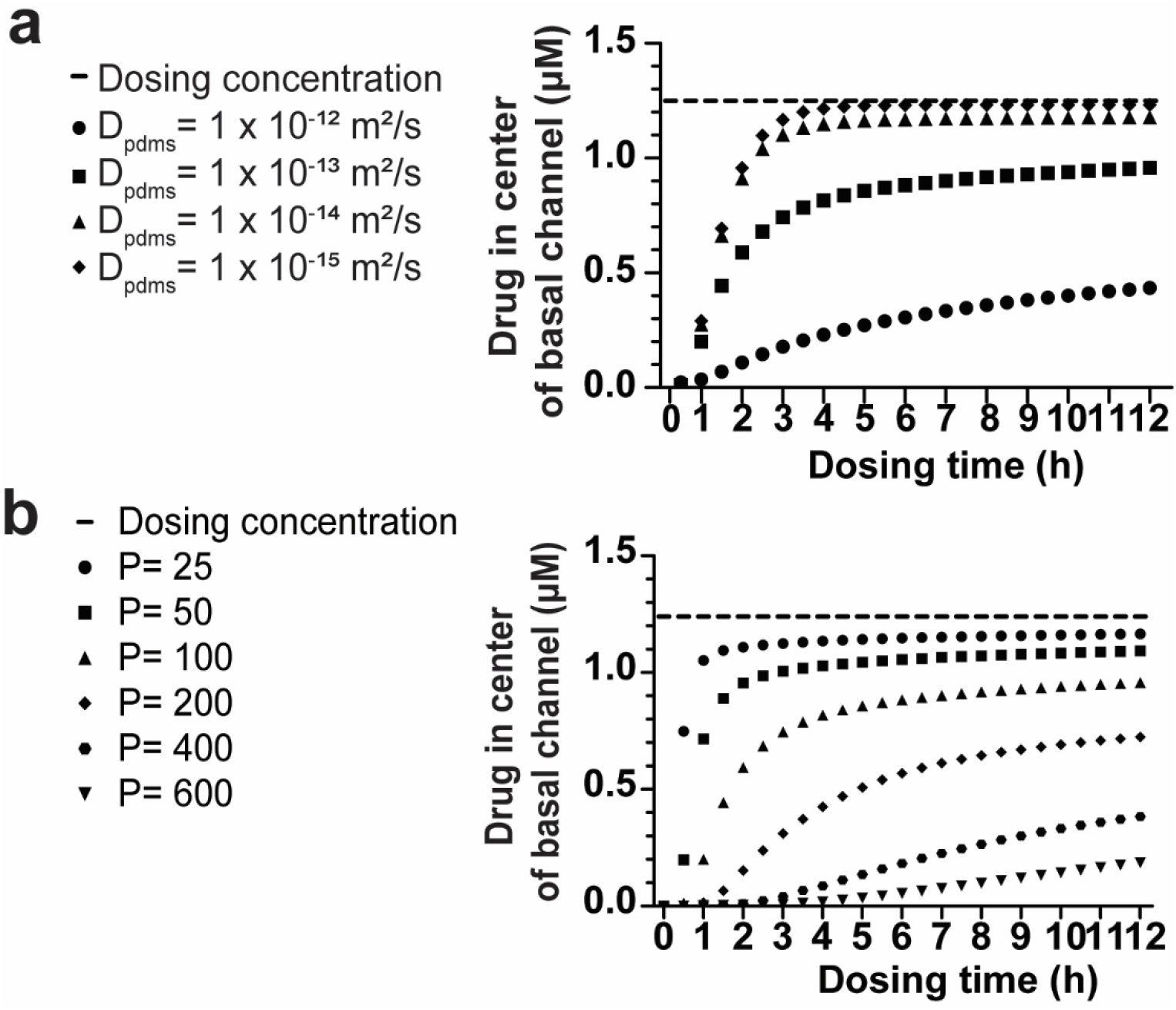
Effect of D_pdms_ and P on the concentration of the drug in the center of the basal channel over time. **a**) Simulated drug concentrations at different D_pdms_ values with P held constant (P=100). **b**) Simulated drug concentrations at different P values with D_pdms_ held constant (D_pdms_= 1 × 10^−13^ m^2^/s).

With the understanding that P and D_pdms_ both contribute to drug loss, we developed a computational and experimental strategy to solve for both values and predict the loss of a drug in the chip over time using amodiaquine (1.24 μM perfused through the basal channel of the human Lung Airway Chip at a flow rate = 60 μL/h) as a test case. Our strategy incorporates three steps: 1) experimentally solve for D_pdms_ of drug, 2) experimentally determine the concentration of drug in the basal outflow, and 3) vary P in the computational simulation to fit the experimental basal outflow concentrations.

### Experimentally solving for the diffusion coefficient

Using amodiaquine to test the approach, the diffusivity of the drug was determined by correlating Fick’s second law to the fluorescence distribution of a surrogate compound inside an Organ Chip. The diffusivity of a compound in a solid polymer is inversely proportional to the molecular size.^26,27^ Fluorescein isothiocyanate (FITC) was selected as a surrogate compound to amodiaquine because it is similar in molecular weight, structure, and hydrophobicity (**Table S1**), and FITC absorption into the bulk PDMS of the Organ Chip can be quantitatively and spatially analyzed using fluorescence microscopy. We flowed FITC into the Organ Chip at 1.24 μM, the C_max_ of amodiaquine, and observed entry into the bulk polymer through the PDMS wall (**Fig. 4a**). The diffusion coefficient was obtained by fitting the analytical solution of Fick’s second law to the experimental data by varying the value of D_pdms_ and minimizing the sum of squares (**Fig. 4b**).^26,28^ We calculated the diffusion coefficient in four regions of the Organ Chip― two regions in the apical channel and two regions in the basal channel― and found that the location in the chip does not affect the value of D_pdms_. The diffusion coefficient that we calculate, D_pdms_ = 3.8 ± 4.5 × 10^−13^ m^2^/s, agrees with previously reported values for compounds with similar molecular weights.^26^

**Figure 4.**
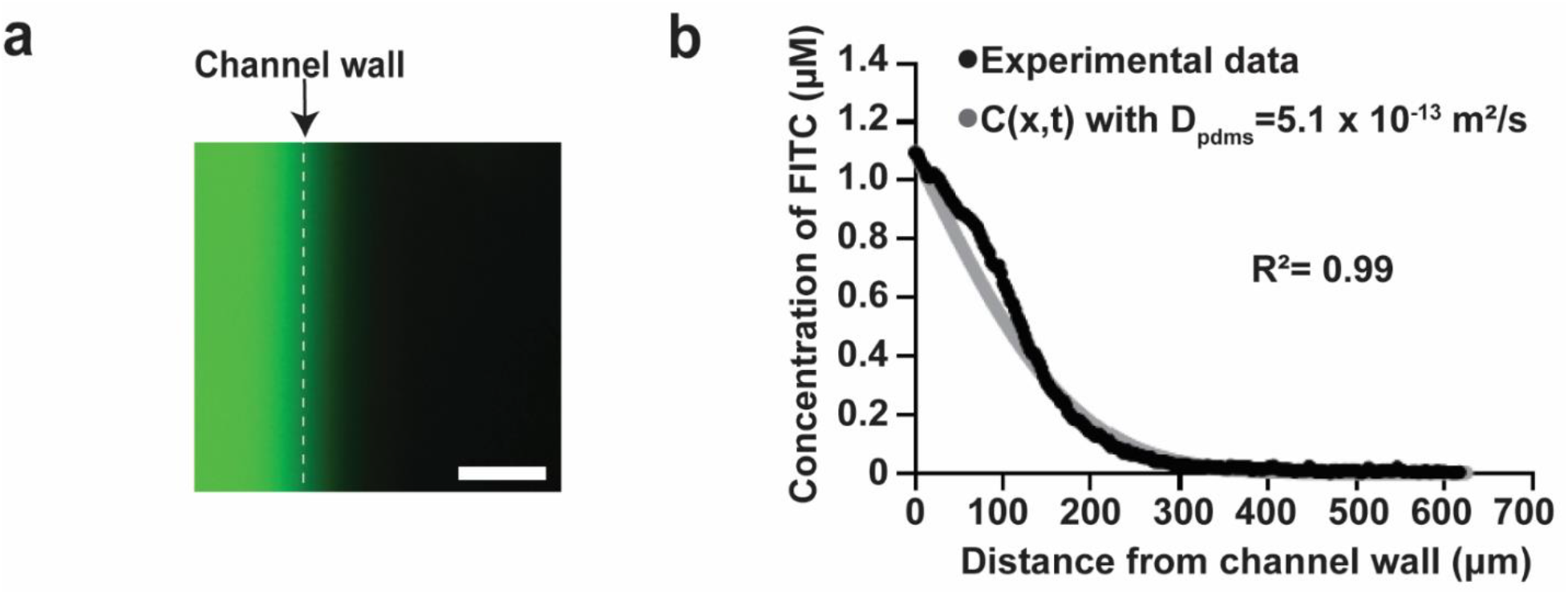
Calculating the diffusion coefficient in PDMS. **a**) Fluorescence microscopic image of FITC in the basal channel (left) at its interface with the channel wall of an Organ Chip after 6 h of continuous flow (bar, 500 μm). Fluorescence is observed extending outward from the surface of the channel wall into the bulk PDMS at the right. **b**) Plot of the normalized fluorescence intensity in bulk PDMS at a distance away from the channel wall. These data, which were acquired from fluorescence images in the center of the apical channel near the device inlet, show that the fluorescence intensity gradually decreases away from the channel wall and the loss is proportional to D_pdms_. The value of D_pdms_ is determined by fitting the analytical solution of Fick’s second law to the experimental data by varying D_pdms_ and minimizing the sum of squares (with R^2^ = 0.99).

### Predicting the Drug Concentration in a Two-Channel Organ Chip

With knowledge of the approximate D_pdms_ value, we estimated P by continuously measuring the concentration of the amodiaquine drug in the outflow over time and fitting the simulation to match the experimental data. Human Lung Airway Chips lined with living bronchial epithelium interfaced with pulmonary microvascular endothelium were perfused with1.24 μM amodiaquine in the vascular channel for 12 h at 60 μL/h and the basal outflow was collected every 3 h. Quantification of amodiaquine in these effluent samples using mass spectrometry reveals that its concentration gradually increases over the dosing period, but does not reach C_max_, because the drug continues to adsorb to the channel walls (**Fig. 5**). We solved for a range of P values that fit the experimental data by simulating the concentration of amodiaquine in the basal channel outflow at steps of P= 5 and terminating the simulation when the concentration exceeded the experimental error at all time points. The simulated basal outflow values fit the experimental data when P= 20-60 (log P= 1.3-1.8). The log P range that we calculate is lower than the log P for amodiaquine reported in DrugBank (log P= 3.7),^29^ because the channels of the human Lung Airway Chip are coated with extracellular matrix and cells that effectively reduce the hydrophobicity and decrease drug adsorption.

**Figure 5.**
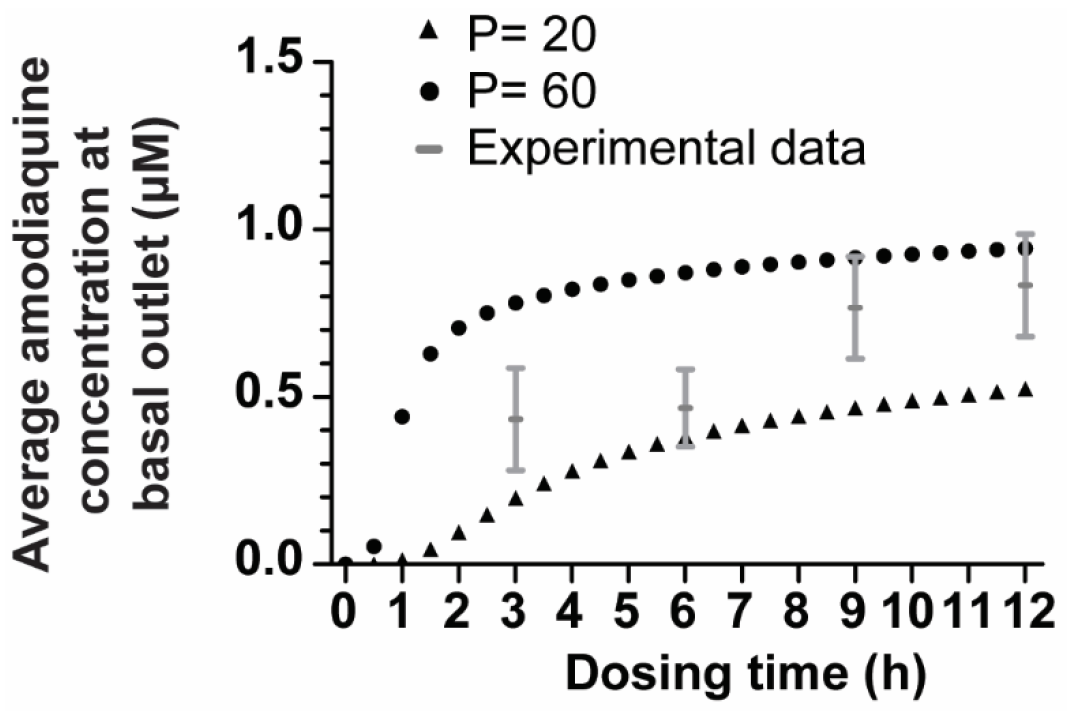
Plot showing the experimentally determined concentrations of amodiaquine in the basal effluent measured over 12 h and predictions that the model fits experimental values for P values between 20 and 60. Simulations were performed using the experimentally determined D_pdms_ value.

Next, we predicted the levels of amodiaquine in human Lung Airway Chips when administered continuously in the basal channel for 72 h used in our past experiment on SARS-CoV-2,^1^ by simulating its concentration in the chip using the calculated values of P= 20 and P= 60 with D_pdms_ = 3.8 ± 4.5 × 10^−13^ m^2^/s. We find that the concentration of amodiaquine inside the basal channel sharply rises during the first 5-10 h of dosing and continues increasing more gradually to reach 0.82-1.08 μM after 72 h (**Fig. 6a**, **Fig. S3**). Heat maps of the amodiaquine concentration profile inside the basal channel at earlier time points show a significant depletion zone near the channel walls (**Fig. 6b, c**). The drug becomes more uniformly distributed across the channel over time because the drug approaches a saturation limit near the PDMS wall. These results reveal that use a pretreatment step would allow the drug concentration to reach near-saturation before introducing a perturbation (e.g. viral or bacterial infection) and that dosing the drug at a higher concentration could compensate for the loss due to adsorption and absorption, and hence allow the *in vitro* model to more accurately approach the clinical C_max_.

**Figure 6.**
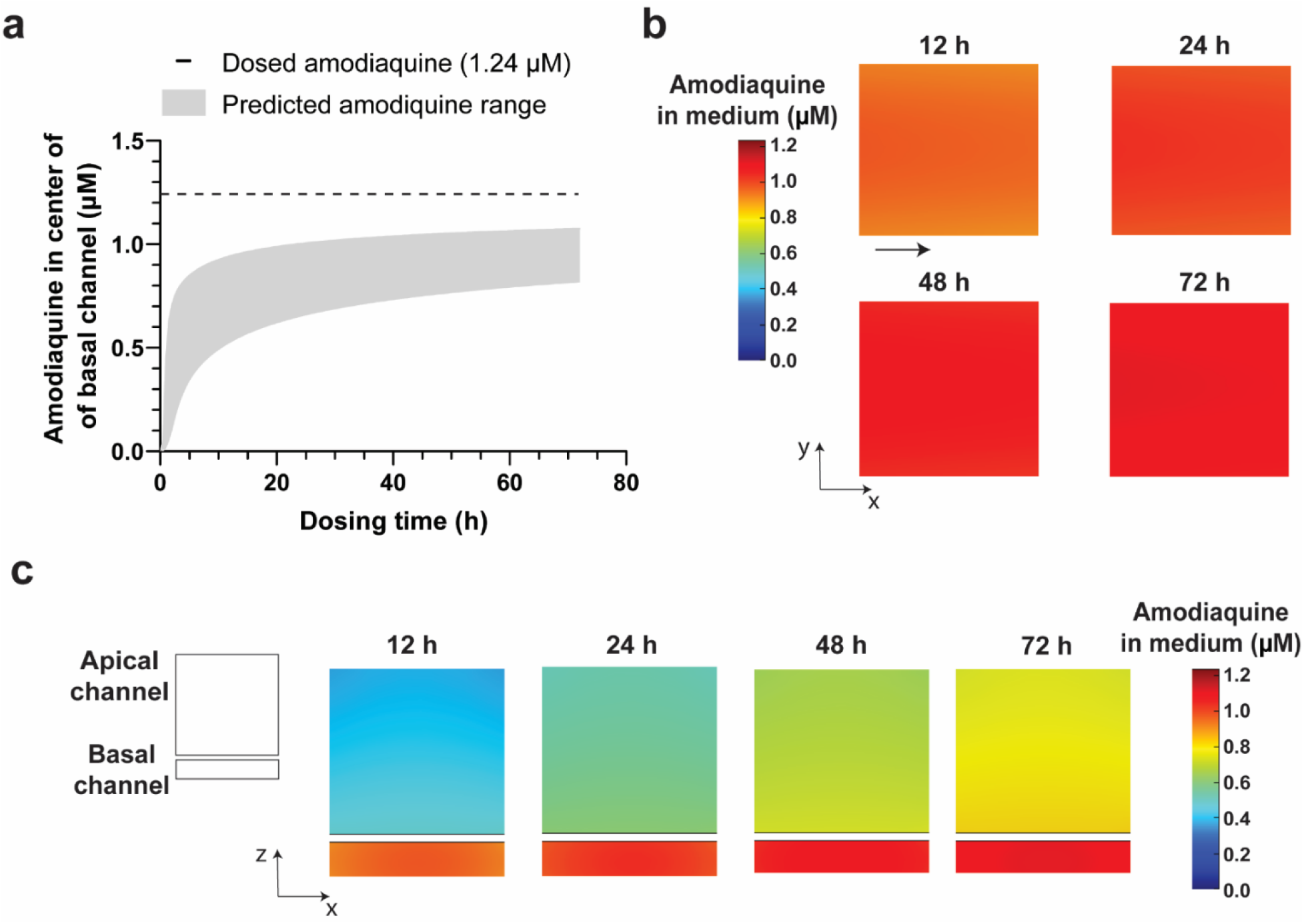
Amodiaquine concentration in the human Lung Airway Chip over a 72 h dosing period. **a**) Plot of the simulated amodiaquine concentration in the center of the basal channel over 72 h at D_pdms_= 3.8 × 10^−13^ m^2^/s and P=20-60 for the predict range (grey). **b**) 2D heat maps of a horizontal cross section through the center of the basal channel along the xy plane at different times with P=40 and D_pdms_= 3.8 × 10^−13^ m^2^/s. A depletion zone is observed near the channel walls at earlier time points. **c**) 2D heat maps of a vertical cross section through center of the chip along the xz plane at different times with P=40 D_pdms_= 3.8 × 10^−13^ m^2^/s showing that the drug evenly fills the lower channel of the chip by approximately 48 h.

## DISCUSSION

The combined experimental-computational strategy described here provides a way to quantitatively predict the spatial and temporal gradient of a drug in 3D within a two-channel PDMS Organ Chip without prior knowledge of the partition coefficient or the rate of diffusion in PDMS. Importantly, this approach addresses a longstanding limitation associated with the use of microfluidic culture devices, which are frequently composed of PDMS, to more accurately inform dosing studies and define toxicity thresholds. We simulated drug loss in 3D and show that it is a spatially and temporally dynamic process that depends on the physicochemical properties of the drug and the experimental Organ Chip model. By experimentally measuring the diffusion coefficient of the compound into PDMS and determining its partition coefficient by measuring its levels in the channel outflow using mass spectrometry, we are able to determine the effective log P range of the compound. We applied this strategy to predict the concentration of amodiaquine in a human Lung Airway Chip after 72 h of continuous dosing and show that the concentration reduces by 13-34% in the center of the basal channel, which affects analysis of prior results and design of future studies with this drug.

Previous work has shown that PDMS can strongly bias drug dosing studies and influence cell behavior. For example, when tested using a multiple myeloma cell line, the IC_50_ for two cancer drugs (bortezomib and carfilzomib) was shown to be 4.3-fold greater in a microfluidic device composed of PDMS compared to polystyrene and cyclo-olefin polymer.^30^ Similarly, human embryonic kidney (HEK) cells responded to fluoxetine (Prozac) treatment when cultured in polystyrene, but not PDMS microchannels.^31^ To demonstrate the effect of PDMS on cell behavior, one study observed a reduction in CD38 expression on human umbilical cord blood-derived CD34^+^ cells cultured on PDMS due to the loss of all-*trans* retinoic acid into the PDMS.^32^ Therefore, understanding and quantifying drug loss in PDMS microfluidic culture systems is critical because it can shift dose-response curves, bias toxicity thresholds, and alter cell behavior.

In a previous study, the loss of drug due to PDMS absorption was incorporated into a PK model of multiple fluidically linked Organ Chips by experimentally measuring the drug in the channel outflows of blank chips and calculating the fractional loss.^10^ In this approach, the geometry of the Organ Chip was simplified into three discrete regions and modeled in 2D, and diffusion of the drug into bulk PDMS was not considered. Our approach advances this work by constructing a 3D model that matches the exact geometry of the Organ Chip and considers all physical processes that contribute to drug loss (e.g., convection, diffusion, adsorption, and absorption). This improved 3D model can predict the concentration of a drug at any location inside the Organ Chip throughout the dosing period. Knowledge of the concentration and distribution of the drug inside the chip allows for the optimization of dosing protocols to ensure that the drug concentration in the chip approaches a clinically relevant dose (e.g., C_max_) before applying a perturbation. Furthermore, we describe a strategy to calculate the effective diffusivity and range of log P values that are specific to a drug under continuous dosing in the *in vitro* model with cultured cells.

Using amodiaquine administration in the human Lung Airway Chip as a test case, we analyzed its distribution in 3D and observed spatial variations in concentration throughout the fluidic channel that varied with time. At earlier time points, a region of drug depletion forms near the channel wall and the concentration decreases along the direction of fluid flow, but over time, the concentration becomes more homogenously distributed throughout the channel. Taken together, these data indicate that drug loss occurs in two phases: the first phase is dominated by adsorption to the channel wall and the second is dominated by diffusion through bulk PDMS. This type of modeling can be useful for retrospective analysis of results of past studies, and to assist in the design of future Organ Chip experiments. For example, our results suggest that a pretreatment step that spans the adsorption-dominating phase would allow a drug to reach near-saturation in the channel before applying a perturbation, and thus, amodiaquine should be flowed through the basal channel of the human Lung Airway Chip for approximately 24 h before infection with the virus in future studies. The analysis also indicates that we might have slightly underestimated the antiviral potency of amodiaquine in our past study^1^ as the inhibition we observed was obtained with a dose that was approximately 25% lower than the clinical C_max_ we originally targeted.

We recognize that individually calculating the D and P values for each compound in a large drug library would be very time-consuming and costly. The computational-experimental approach described here can be used in the future to refine existing QSPR^33,34^ and QSAR^35^ models and permit *in silico* predictions of the diffusion and partition coefficients of a drug in the Organ Chip. QSPR and QSAR models use descriptors that represent the molecular structure and properties of a compound to predict an unknown activity or property,^36–38^ and experimental determination of D and P values for a training set of compounds can help to refine QSPR and QSAR models, respectively. For example, the diffusivity of a training set of diverse fluorescent compounds may be experimentally determined and used to build a QSPR model that is predictive of the experimental data. The descriptors in the QSPR model can then be iterated to maximize the correlation between experimental and predicted diffusivities. This strategy would circumvent the need to use a fluorescent surrogate compound because the diffusivity of the actual drug will be derived from the QSPR model. Similarly, a QSAR model can be iterated using P values derived from our computational-experimental approach for a representative group of molecules. We note that mass spectrometric analysis of the drug concentration in the outflow of the Organ Chip should still be performed for each drug and chip design to verify that the simulated outflow concentration matches the actual concentration present in the medium.

It is important to note that another potential limitation of our approach is that the PDMS membrane, epithelium, and endothelium are modeled as a homogenous PDMS block, and we assume that drug passively diffuses through the block at a rate described by Fick’s second law. The accuracy of the predicted range of log P values may be improved in the future by measuring accumulation of the drug in the cells and apparent drug permeability (P_app_), and incorporating this information into the model. The concentration of drug in the basal channel outflow can then be combined with the simulation of drug loss and fit to a PK model to estimate the rate and extent of distribution.^19^ Furthermore, P_app_ can be determined by flowing a tracer molecule into the basal channel and calculating the fraction of molecule that entered the apical channel.^1,10,39^

Organ Chips and other microfluidic culture models are commonly fabricated with PDMS, but compound loss can bias results and affect cell behavior. In this study, we described a combined experimental and computational strategy to predict the concentration of a drug inside a two-channel microfluidic Organ Chip composed of PDMS under continuous vascular dosing that considers convection, diffusion, and dissolution of the drug in the chip. Using this strategy, drug concentration can be estimated over a multi-day dosing period without prior knowledge of the partition or diffusion coefficient. The 3D computational simulations of drug loss emphasize that an understanding of spatial and temporal drug loss is necessary to establish dosing guidelines. In addition, a single coefficient derived from the literature or a chemical database is likely not applicable for an *in vitro* microfluidic model because drug loss depends on the relationship between the physicochemical properties of the drug, the duration of dosing, and the design of the Organ Chip device used for therapeutic testing. Our strategy may be used in the future to characterize drug loss for a range of compounds and microfluidic PDMS devices. It also may increase the usefulness of Organ Chips in clinical research programs by offering a way to more accurately plan dosing strategies and predict drug pharmacokinetics, therapeutic windows, and toxicities in a human organ-relevant context *in vitro*.

## METHODS

### Computational simulations

A finite element simulation of drug concentration in the human Lung Airway Chip was developed in COMSOL Multiphysics 5.5 (COMSOL, Inc.). A SOLIDWORKS (Solidworks 2017) 3D rendering of the microfluidic device was imported into COMSOL. The Organ Chip is a (PDMS) microfluidic culture device (37.2 mm long × 16.2 mm wide) that contains parallel fluidic channels (15.5 mm long × 1 mm wide) separated by a porous PDMS membrane (50 μm thick; 7 μm pores). The fluidic component consists of two adjacent parallel microchannels (apical, 1 mm wide × 1 mm high; basal, 1 mm wide × 0.2 mm high; length of overlapping channels, 16.7 mm). The height of the epithelium (3.9 μm) and endothelium (1.0 μm) was experimentally determined from cross-sections of human Airway Chips. The epithelium and endothelium layers were combined with the membrane to create a single geometric domain. The membrane pores were removed to increase computational efficiency.

The drug flow was simulated using the Navier-Stokes equations for incompressible flow coupled with continuity law, and assuming incompressible flow and constant viscosity across the domain:

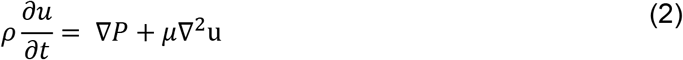

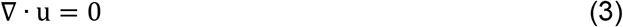

where is ∇ the gradient operator and ∇^2^ is the square of vector Laplacian, P is pressure, u is velocity, *μ* is viscosity, and *ρ* is the density of medium. Apical and basal inlet flow rates were set as 0 μL/h and 60 μL/h, respectively, for the flow simulations. The outlet boundary condition was set to zero gauge pressure and a no-slip boundary condition was set at the channel walls.

Fick’s second law was applied to model drug transport in culture medium (Equation #4) and PDMS (Equation #5):

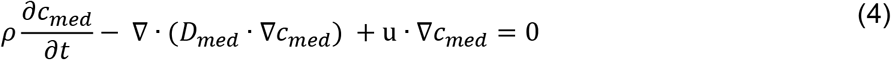

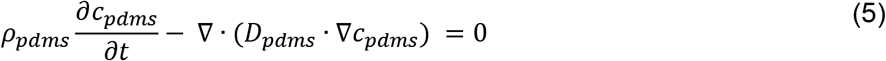

where *D*_*med*_ is the diffusivity of drug in cell culture medium, *D*_*pdms*_ is the diffusivity of drug in PDMS, *c*_*med*_ is the concentration of drug in medium, *c*_*pdms*_ is the concentration of drug in PDMS, and *ρ*_*pdms*_ is the density of PDMS.

The partition coefficient was implemented in COMSOL by applying the following pointwise constraint expression at all PDMS-medium and PDMS-cell interfaces:

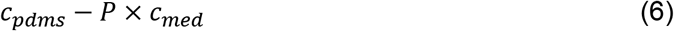

where *test*(*c*_*pdms*_) − *test*(*c*_*med*_) was implemented in the constraint force expression field to ensure continuous flux over the phase boundaries. A zero flux boundary condition was applied on the walls and ports of the vacuum channels. A custom meshing sequence was carried out in order to increase the mesh resolution near PDMS-medium boundaries, within the fluidic domain, and near corners of the fluidic domain. The laminar flow and diffusion of dilute species modules were coupled and a time-dependent simulation was performed for 0-12 h with 0.5 h time steps or for 0-72 h with 0.5 time steps. The concentration of drug in the basal outflow was calculated by placing a boundary probe on the basal outlet and programming the probe to report the average drug concentration. Similarly, the concentration of drug in the center of the basal channel was determined by placing a point probe in the middle of the basal channel and programming the probe to output the concentration of the drug. We also simulated the concentration in the basal channel without dosing 1.24 μM statically in the apical channel and observed no effect on the concentration of drug. Parameters used in the simulation are provided in **Table S2**.

### Cell Culture

This protocol is based on the methods described in Si et al. with minor modifications.^1^ Microfluidic Organ Chips were purchased from Emulate, Inc (CHIP-S1 Stretchable Chip, RE00001024 Basic Research Kit). The Organ Chips have two fluidic microchannels that are separated by a membrane containing 7 μm diameter pores. To prepare the chips for cell culture, the surface of the fluidic channels and membrane were activated with ER1+ER2 (Emulate, Inc) in the presence of UV light (Nailstar, NS-01-US) for 20 minutes. The channels were washed with ER2 buffer, followed by PBS −/−. The activated fluidic channels were coated with 0.5 mg/mL collagen type IV human placenta (Sigma Aldrich, C5533-5MG) in the cell culture incubator overnight at 37 °C, 100% humidity and 5% CO_2_. The coating solution was aspirated from the chip the following day.

Human lung microvascular endothelial cells (HMVEC-L) were sourced from LONZA (CC-2427) and cultured in EGM-MV2 medium supplemented with growth factors and 1% penicillin-streptomycin (Gibco,15140122) according to the manufacturer’s instructions (EGM-MV2 Kit, Promocell, C-22121). Endothelial cells were seeded onto the top and bottom of the basal channel at 2 × 10^7^ cells/mL, passage 2 (p2), and incubated statically for 2 h at 37 °C. Primary normal human bronchial epithelial (NHBE) cells were sourced from LONZA (CC-2540). The epithelial cells were cultured and expanded in Airway Growth Medium (Airway Epithelial Growth Medium Kit, Promocell, C-21160) supplemented with 1% penicillin-streptomycin. NHBEs were seeded into the apical channel of the Organ Chips at a density of 2.5 × 10^6^ cells/mL at passage 2 (p2) and incubated for 4 h under static conditions at 37 °C.

The next day, the chips were inserted into Pod Portable Modules (RE00001024 Basic Research Kit; Emulate, Inc) and placed in a Zoë Culture Module (Emulate, Inc). EGM-MV2 and Airway Growth Medium was added to the basal and apical inlet reservoirs, respectively. The Zoë was programmed to flow at 30 μL/h medium into the apical and basal channel and the chips were maintained cells were maintained at 37 °C, 100% humidity and 5% CO_2_. Air-Liquid Interface (ALI) was established after 5-7 days. After establishment of ALI, the apical channel medium was removed from the human Lung Airway Chip and the basal medium was replaced with PneumaCult-ALI medium (PneumaCult™-ALI Medium, StemCell Technologies, 05001) supplemented with 1% penicillin-streptomycin, 10 ng/mL of VEGF and 1 μg/mL ascorbic acid. The apical channel of the chips were flushed for 2 minutes at 1000 μL/h. After flushing, the flow rate in the apical channel was set to 0 μL/h and the Zoë was programmed to “Air”. Medium flowed through the basal channel at 30 μL/h. The apical surface of the epithelium was rinsed once weekly with PBS (−/−) to remove cellular debris and mucus. Drug was dosed into human Lung Airway Chips after 12 days of ALI culture.

### Diffusion Coefficient (D_pdms_) Determination

A blank Organ Chip (CHIP-S1 Stretchable Chip, RE00001024 Basic Research Kit; Emulate, Inc) was equilibrated with PBS (-/-) and placed inside a microscope chamber at 37 °C and 100% humidity to match the environment of a cell culture incubator. A 1.24 μM solution of FITC (Sigma Cat# 46950) in PBS was introduced into the apical and basal channel at 60 μL/h for 7 h with a syringe pump (Harvard Apparatus). Four regions of the chip (two in the apical channel and two in the basal channel) were imaged after 6 h of flow at 10 × magnification (Zeiss Axio Observer Z1.2). The fluorescence intensity across the microfluidic channel was quantified using ImageJ and normalized to the dosing concentration. The diffusion coefficient of FITC in PDMS was determined by applying the analytical solution of Fick’s second law:

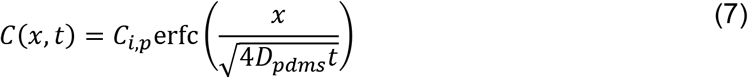

where *C*_*i,p*_ is the concentration of the drug on the PDMS side of the medium-PDMS interface, *x* is the distance away from the wall into bulk PDMS, *D*_*pdms*_ is the diffusion coefficient of the drug in PDMS, *t* is the elapsed time, and *C*(*x, t*) is the concentration of the compound. *C*(*x, t*) was fit to the experimentally determined concentration profile by varying D_pdms_ and minimizing the sum of squared residuals.^26,28^ The diffusion coefficient was calculated from n=3 experimental replicates and R^2^> 0.98 for each replicate.

### Amodiaquine Dosing

Amodiaquine dihydrochloride dihydrate (cat. #A2799; Sigma-Aldrich) was dissolved in dimethyl sulfoxide (DMSO) to a stock concentration of 10 mM and diluted in ALI medium to the reported C_max_ in human blood (1.24 μM). Amodiaquine was perfused through the vascular channel of human Lung Airway Chips at 60 μL/h while the airway channel was statically treated with the same concentration of drug. Medium from the inlet and outlet of the vascular channel was collected every 3 h for 12 h and frozen at −80 °C until LC-MS/MS analysis. Mass spectrometry was performed by PureHoney Technologies (Billerica, MA) using a RapidFire-MS/MS system. The RapidFire-MS/MS system consisted of an Agilent RapidFire 300 interfaced to a Sciex API4000 triple quadrupole mass spectrometer with Analyst v.1.6 data acquisition software. Quantitative analysis was performed with Agilent RapidFire Integrator software. Standard curves were generated with 0-4 μM amodiaquine and all samples were run in triplicate. The concentration of amodiaquine in the basal channel outflow was normalized to the concentration in the basal inlet to account for adsorption of the drug to the Pod walls that occurs within the first 6 hours of dosing.

### Partition Coefficient (P) Determination

The concentration of amodiaquine in the basal outflow was computationally simulated using the experimentally determined D_pdms_ 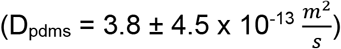. A boundary probe was placed on the basal outlet and programmed to output the average drug concentration. We first performed a course estimate of the P value range that fit the LC-MS/MS data. We performed a time-dependent simulation for 12 h with 0.5 h time steps at P= 25, 50, 100, and 200 and found that the basal outflow concentrations at P~15-100 approximate the LC-MS/MS data. We solved for the range of P that fit the experimental data by simulating the concentration of amodiaquine at steps of P= 5 and terminating the simulation when the concentration exceeded the experimental error at all time points. We determined the lower and upper boundary of P by initiating the stepwise simulation at P=15 and P= 100, respectively. P= 20-60 fit the experimental LC-MS/MS data (log P= 1.3-1.8).

### Statistical analysis

All experimental results are expressed as means ± standard deviation (SD). Each experiment was conducted with a sample size of n=3 chips per condition.

## ACKNOWLEDGMENTS

This work was supported by DARPA under Cooperative Agreements (W911NF-16-C-0050 and HR0011-19-2-0008/AWD-102840-612, NIH (NCATS 1-UH3-HL-141797-01 to D.E.I.), and the Wyss Institute for Biologically Inspired Engineering at Harvard University.

## CONTRIBUTIONS

J.G. and D.E.I. conceived this study. J.G., A.O., and C.O. performed and analyzed experiments and simulations with other authors assisting with data analysis. R.P. designed and executed drug dosing experiments. R.P. and D.E.I. managed the project progress. J.G. and D.E.I. wrote the manuscript with all authors providing feedback.

