## Supplementary Information for "Predicting drug concentrations in PDMS microfluidic organ chips"

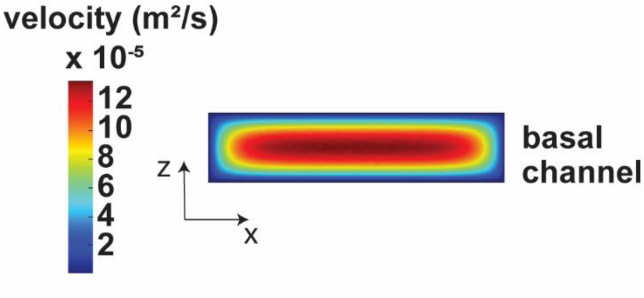


**Figure S1.** Heat map of the velocity distribution in a vertical cross section of the basal channel along the xz axis.

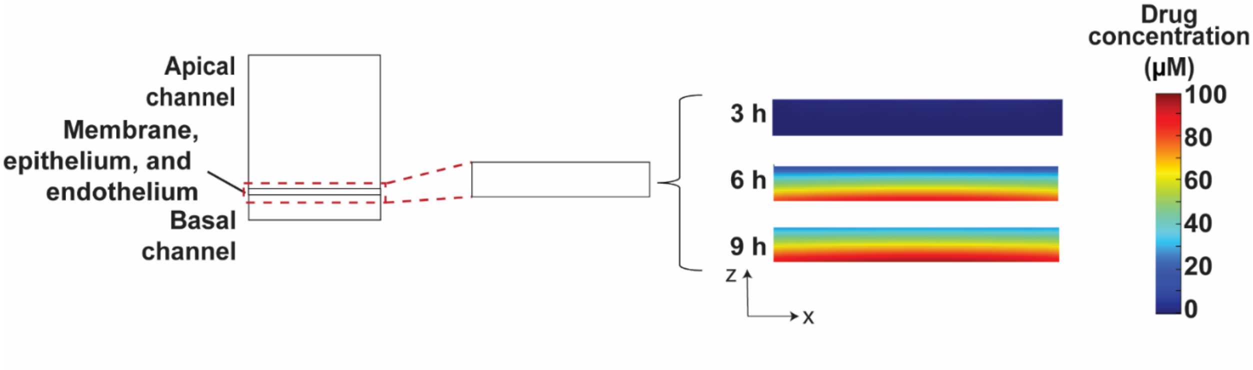


**Figure S2**. 2D heat maps of the concentration profiles of the drug in vertical cross sections of the epithelium, membrane, and endothelium along the xz axis. The epithelium, membrane, and endothelium are combined into one geometry.

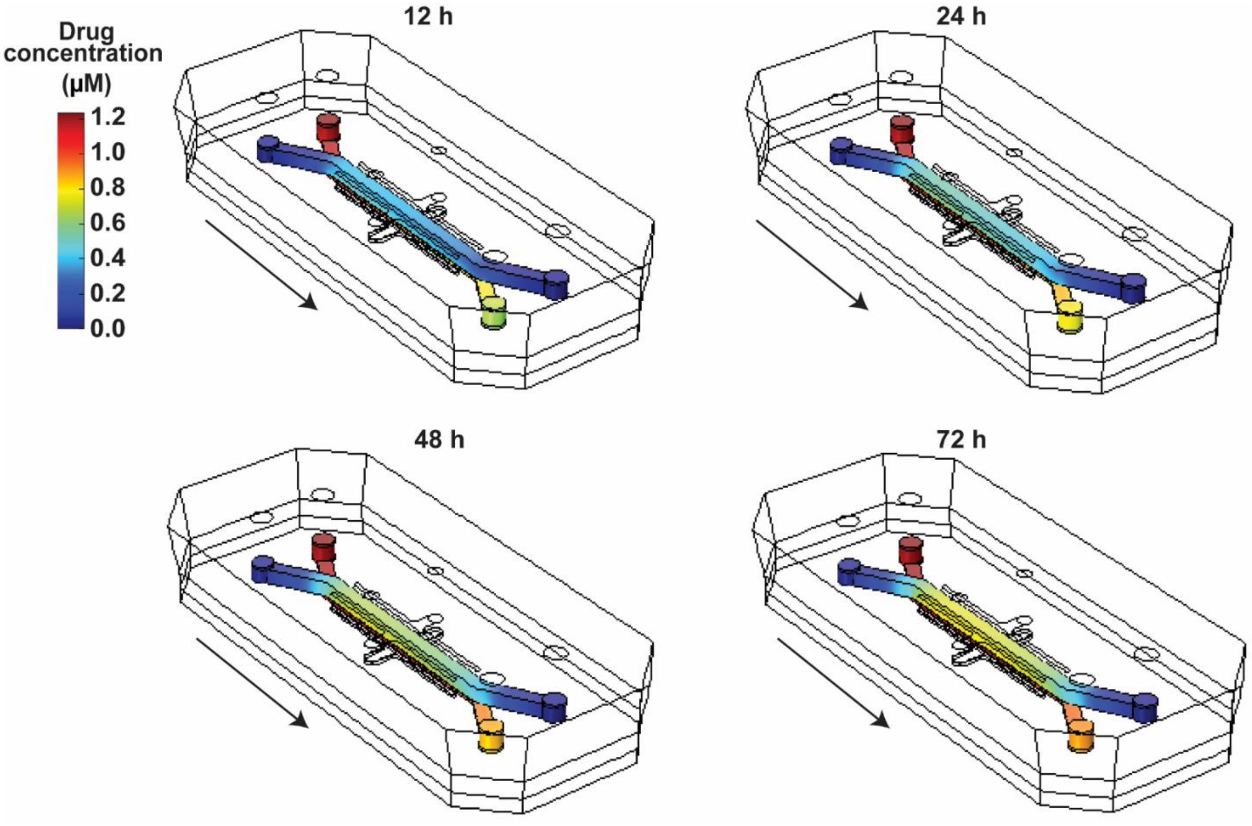


**Figure S3.** 3D surface heat maps showing amodiaquine concentrations in the Organ Chip over a 72 h dosing period. The heat maps were generated using P= 40 and $D_{pdms}$ = 3.8 x $10^{-13}$ $\frac{m^2}{s}$.

**Table S1. Physiochemical properties of amodiaquine and FITC**.

| | **Amodiaquine (from DrugBank)**[1] | **FITC (from PubChem)**[2] |
|---|---|---|
| **Molecular weight (g/mol)** | 355.86 | 398.4 |
| **Log P** | 3.7 | 4.8 (XLogP2 value) |
| **Hydrogen acceptor count** | 4 | 7 |
| **Hydrogen donor count** | 2 | 2 |
| **Number of rings** | 3 | 5 |

**Table S2.** Model parameters.

| Parameter | Description | Value | Unit | Ref |
|---|---|---|---|---|
| $\boldsymbol{Q}$ | Basal channel flow rate | 60 | $\frac{\mu L}{h}$ | |
| $\boldsymbol{T}$ | Incubator temperature | 37 | ℃ | |
| ***P*** | Atmospheric pressure | 1 | *atm* | |
| $\boldsymbol{D_{med}}$ | Diffusion coefficient of the drug in medium | $1 \times 10^{-9}$ | $\frac{m^2}{s}$ | 3–5 |
| $\boldsymbol{c_o}$ | Concentration of drug dosed into the chip | 1.24 | *µM* | |
| $\boldsymbol{D_{pdms}}$ | Diffusion coefficient of the drug in PDMS | Unknown | $\frac{m^2}{s}$ | |
| $\boldsymbol{P}$ | Partition coefficient of the compound | Unknown | | |
| $\boldsymbol{c_{pdms}}$ | Concentration of the drug in PDMS | Unknown | *µM* | |
| $\boldsymbol{c_{med}}$ | Concentration of the drug in cell culture medium | Unknown | *µM* | |
